# Thymidylate synthase disruption to limit cell proliferation in cell therapies

**DOI:** 10.1101/2023.06.26.546441

**Authors:** Rocio Sartori-Maldonado, Hossam Montaser, Inkeri Soppa, Solja Eurola, Melanie Balaz, Henri Puttonen, Timo Otonkoski, Jonna Saarimäki-Vire, Kirmo Wartiovaara

**Affiliations:** Stem Cells and Metabolism Research Program, Faculty of Medicine, University of Helsinki, Helsinki, Finland; Children’s Hospital, University of Helsinki and Helsinki University Hospital, Helsinki, Finland; Clinical Genetics, Helsinki University Hospital, Helsinki, Finland; Department of Pathology, Helsinki University Hospital, Helsinki, Finland

## Abstract

Engineered cells hold great promise for regenerative medicine and gene therapy. However, living cell products entail a fundamental biological risk of unwanted growth. Here, we describe a novel metabolic safety system to control cell proliferation without added genetic elements. We inactivated a key enzyme for nucleotide metabolism, *TYMS*, in several cell lines, thus obtaining cells that proliferate only when supplemented with exogenous thymidine but fail to replicate in its absence. Under supplementation, *TYMS*^-/-^ pluripotent stem cells proliferate normally, produce teratomas and differentiate into potentially therapeutic cell types such as pancreatic beta cells. After differentiation, the postmitotic cells do not require thymidine to function, as seen by prolonged *in vivo* production of human insulin in implanted mice. Hence, this method allows robust cell culture and manufacture while mitigating the risk of uncontrolled growth of transplanted cells.

**One Sentence Summary:** Genetic disruption of DNA synthesis prevents unwanted proliferation in cell therapies without affecting cell function.

## Introduction

Somatic cell therapies offer new possibilities for previously untreatable or challenging medical conditions. Among the cell types used in such therapies, induced pluripotent stem cells (hiPSCs) have garnered significant attention due to their availability from patient-specific somatic cells (*1*, *2*) and their capability to differentiate into potentially therapeutic cell lineages, such as glucose-responsive insulin secreting pancreatic islets (*3*, *4*). Furthermore, the improved technologies for gene transfer and genome editing enable the development of genetically modified cell-based therapies with enhanced therapeutic potential (*5–7*).

The use of hiPSC offers several advantages over mesenchymal and embryonic stem cells (MSC and ESC, respectively). HiPSCs circumvent ethical concerns associated with the embryo derived ESCs (*8*, *9*). Unlike donor-derived MSCs and ESCs, hiPSCs’ patient-specificity minimizes the risk of immune rejection and graft-versus-host disease (*10*, *11*). Moreover, stem cells can be further genetically modified by silencing or modifying genes involved *e.g.* in immune recognition (*12*). These features broaden their applications in disease modeling and therapy development, and may enhance their compatibility for allogeneic cell therapies, reducing the need for immunosuppression compared to other cell sources. However, although the universal off-the-shelf cell products decrease the risk for immune rejection, this diminished immunogenicity can also lead to problematic immune evasion and potential oncogenic events.

On top of creating biologically useful cells for a desired clinical purpose, a successful therapeutic product needs to meet the criteria for safety and manufacture scalability. These two goals, however, often compromise each other, since manufacturing benefits from robust cell growth, but intensive or unlimited proliferation poses a threat when the cells are transferred to patients (*13*, *14*). Attempts to create safer cells for therapy have yielded different strategies (*15*, *16*). These safety systems aim to prevent uncontrolled proliferation or selectively eliminate the transplanted cells if necessary (*17*, *18*). Some of the existing safety strategies include the use of inducible apoptotic systems, suicide genes, antibody-mediated cell depletion, and gene editing-based switches (Table 1). Often, these switches work by reversibly activating or inactivating genes upon exposure to a small molecule (ON- and OFF-switches, respectively). However, they all present remarkable challenges: susceptibility to genetic silencing or alteration, transgenic or viral-derived origin, slow activation speed, rapid remission and/or potential neurotoxicity (*16*, *19*, *20*).

In this context, we envisioned the generation of an *endogenous* safety strategy by disrupting a key metabolic gene, whose function can be compensated by administering an easily available compound. Using CRISPR-Cas9, we genetically inactivated thymidylate synthase (*TYMS*), the only known enzyme in charge of the *de novo* thymidylate (dTMP) synthesis (*21*). This knock-out generates rescuable auxotrophy towards thymidine and restricts the synthesis of DNA, without altering normal RNA or protein production (Fig. 1.A). This means that dividing cells depend on external supplementation of thymidine to proliferate and achieve robust manufacturing of the cells of interest. However, once the cells have exited the cell cycle and terminally differentiated, they do not require supplementation to synthesize nucleic acids. While preventing dividing cells from proliferating uncontrollably, this approach does not require the use of small molecules nor the insertion of large transgenic elements. Hence, it cannot be silenced by mutagenesis, and it does not result leaky or immunogenic.

**Figure 1.**
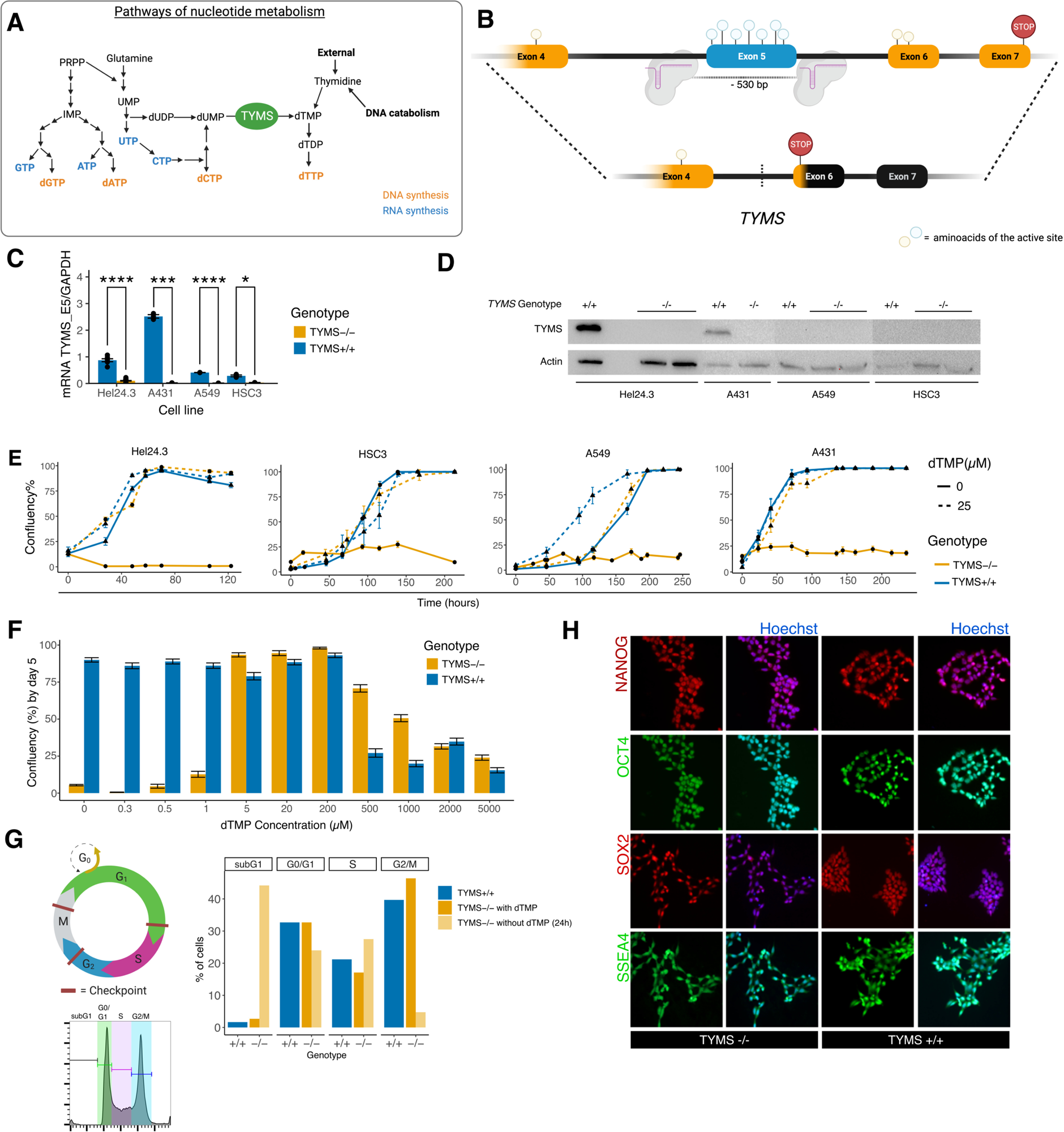
**A.** Diagram showing the nucleotide pathways for DNA (orange) or RNA (blue) synthesis. **B.** Schematic representation of the gene of interest *(TYMS),* showing the two gRNAs, and the amino acids that form the active site of the protein over the corresponding exon that code for them. The lower line represents the change of frame upon DNA repair. **C.** TYMS expression in wild-type versus knock-out hiPSC (HEL24.3), A431, A549 and HSC3 (n=3/cell line). Results shown as ratio of *TYMS* mRNA (primer targeting exon 5) over *GAPOH* mRNA contents. **D.** Western blot showing the absence of TYMS upon inactivation of its gene. **E.** Growth curve of wild-type and knock-out cells supplemented with O or 25 µM dTMP. Results shown as average confluency ± SE. **F.** Confluency at day 5 of wild-type and knock-out hiPSC under different concentrations of dTMP (0 to 5000 µM). Results shown as average confluency per image ± SE. **G.** hiPSC immunocytochemistry analysis of pluripotency markers NANOG, OCT4, SOX2 and SSEA4, alone or merged with a nuclear dye channel (Hoechst). **H.** Summary of flow cyctometry results for cell cycle analysis.

Our results provide evidence that the culture and proliferation of *TYMS*^-/-^ cells can be regulated with exogenous dTMP *in vitro* and *in vivo*. Furthermore, the characterization and differentiation of *TYMS*^-/-^ -hiPSC show that they provide a functional source for cell therapy that does not sustain unwanted proliferation without external dTMP supplementation.

## Results

### Disruption of *TYMS* makes proliferative cells dependent on thymidine supplementation

The human thymidylate synthase (PDB: 5X5D) is evolutionarily highly conserved. The functional enzymatic ligand pocket is encoded by exons 4 (amino acid R175), 5 (P193, C195, Q214, S216, D218, R225) and 6 (H256, Y258) (*22*). Therefore, to disrupt TYMS activity, we deleted the exon 5 with a dual sgRNA Cas9 knock-out approach targeting both intron 4 and intron 5. (Fig. 1.B). This removed the enzymés active site, generated an early stop codon in exon 6 and depleted the enzyme expression at both mRNA and protein levels (Fig. 1.C-D, Fig. S1.A-B) in several immortal cell lines.

We hypothesized that *TYMS^-/-^* cells could be cultured and manufactured normally with external dTMP supplementation, but they would cease to proliferate in its absence. Therefore, we followed the growth of *TYMS*^-/-^ -hiPSC (HEL24.3) (*23*) and cancer cell lines without or with dTMP supplementation (25 µM). The untreated (*TYMS*^+/+^) cell lines were used as controls. The results suggest that *TYMS^-/-^* cells proliferate at a rate comparable to *TYMS*^+/+^ when supplemented with thymidine, while the proliferation capacity of non-supplemented *TYMS^-/-^* cells was drastically impaired (Fig. 1.E). To determine the optimal concentration of dTMP for culturing TYMS^-/-^ -hiPSC, we run a dose-dependent assay using a range of 0.3 uM to 5 mM exogenous dTMP to explore their tolerance limits (Fig. 1.F and Fig. S1.C). Contrarily to *TYMS*^+/+^ lines, *TYMS*^-/-^ -hiPSCs could not sustain growth at physiological concentrations of thymidine (reference: 0.5-1.4 µM) (*24*). Cell cycle analysis by flow cytometry of *TYMS*^-/-^ -hiPSC cells reveal that in the first 24 h of dTMP withdrawal, the cells accumulate in G0-G1 and S-phase, and then, as they should continue to G2, accumulate in the subG0-G1 phase (Fig. 1G and Fig. S1.D). On the other end of this spectrum, we observe shift in the cytotoxic threshold at higher concentrations; while the growth rate of *TYMS*^+/+^-cells decreases at concentrations higher than 200 µM, knock-out cells suffer comparable cytotoxic effects at concentrations higher than 1 mM (1000 µM) (Fig. 1.F).

In summary, our results show that *TYMS*^-/-^ cells grow normally and can be expanded extensively under dTMP supplementation. Thymidine withdrawal, however, stalls cells in S phase and inhibits progression to G2/M, likely due to replicative stress from nucleotide imbalance.

### *TYMS* knock-out hiPSCs maintain pluripotency and genomic integrity

The *TYMS*^-/-^ -hiPS cell lines performed equally well on standard pluripotency analysis as their non-edited counterparts. They showed positive staining for pluripotency markers NANOG, OCT4, SOX2 and SSEA4 and comparable mRNA expression of *SOX2, NANOG* and *OCT4* to *TYMS*^+/+^-hiPSC (Fig. 1.G and Fig. S1.E). Furthermore, their chromosomal integrity remained unaltered by the editing process in prolonged culture, tested in several time points for up to 25 passages (Fig. S1.F). The differentiation into the three germ layers with or without dTMP supplementation revealed that the cells successfully differentiate into ectoderm, mesoderm and endoderm when supplemented with dTMP at the early stages of differentiation (Fig. 2.A, C). Lack of supplementation from the *TYMS*^-/-^ cells during d0-d14 produced significantly less embryoid bodies at d14 (Fig. 2.B). By day 28, however, all attached cells from supplemented and non-supplemented cells expressed characteristic markers of the three germ layers (Fig 2.C, Fig. S1.G).

**Figure 2.**
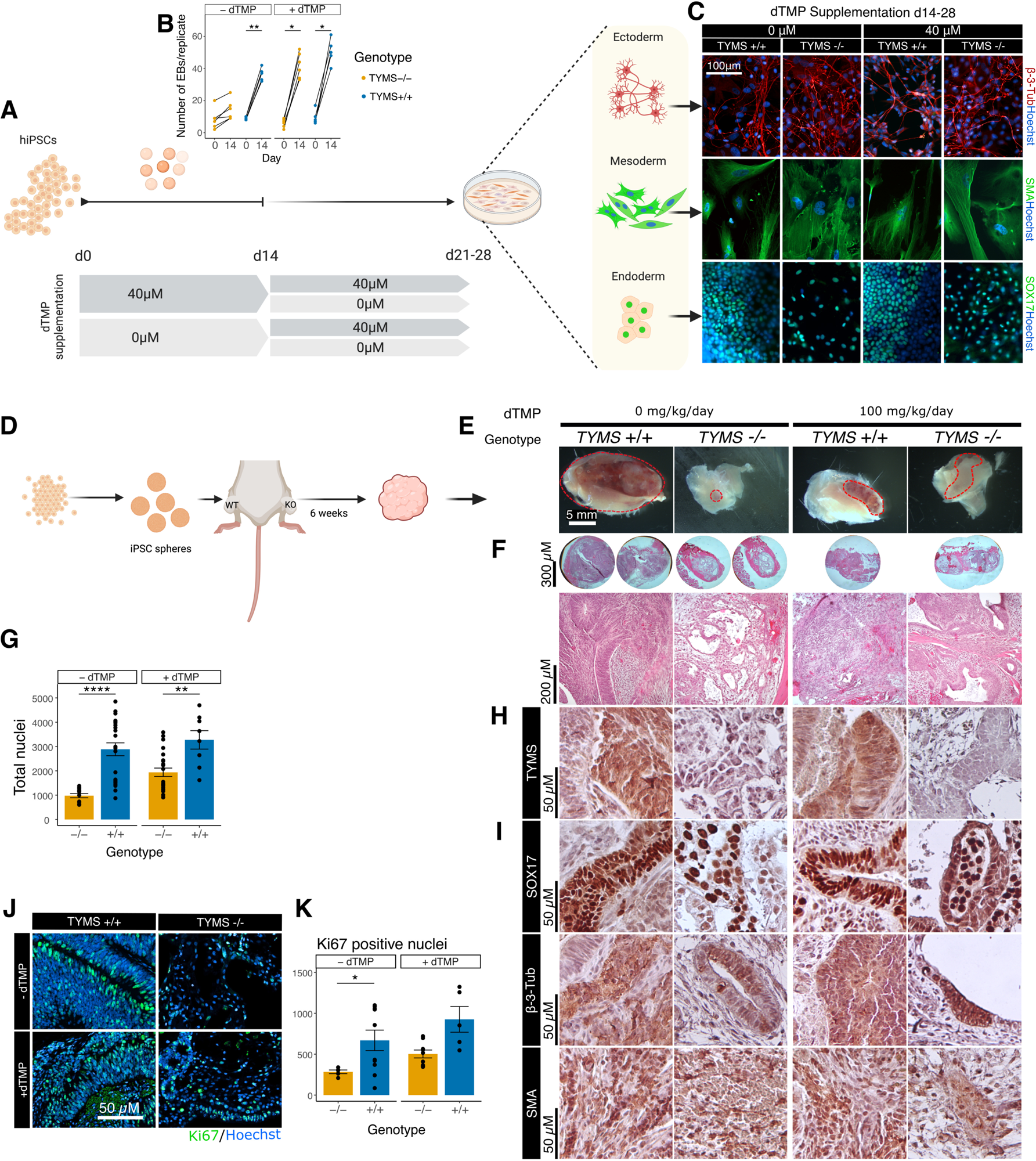
**A.** Graphical representation of trilineage differentiation protocol. **B.** Number of resulting 300 *µm* aggregates from wild-type and knock-out hiPSC with and without supplementation during the first stage of differentiation (day O to 14). **C.** lmmunocytochemisty analysis of markers for ectoderm (beta-3-tubulin), mesoderm (smooth muscle actin, SMA), and endoderm (SOX17) in cells derived from wild-type and knock-out hiPSC under dTMP supplementation during the first stage of differentiation. **D.** Graphical representation of teratoma formation experiment. **E.** Knock-out and wild-type hiPSC-derived teratomas with and without dTMP supplementation as extracted from the mice, before processing for molecular analysis. **F.** Macro and micro images of haematoxylin-eosin staining show dense and cystic structures within the tumors. **G.** Quantification of cellularity (cell density), based on nuclear count of sectioned teratomas. **H.** lmmunohistochemistry against TYMS. I. lmmunohistochemistry of characteristic markers of endoderm (SOX17), ectoderm (beta-3-tubulin) and mesoderm (SMA). **J.** lmmunohistochemistry against Kl67 (proliferation marker, green) and total nuclei (Hoechst, blue). **K.** Quantification of Kl67 positive nuclei from immunohistochemistry. Statistical significance in panels B, G and K based on Wilcoxon test; p>0.05 (ns, not shown), p < 0.05 (*), p < 0.01 (**), p < 0.001 (***), p < 0.0001 (****).

### Thymidine deficiency drastically reduces proliferation and teratoma formation in mice transplanted with TYMS^-/-^ -hiPSCs

Next, we aimed to translate the *in vitro* trilineage differentiation to an *in vivo* setting. We implanted undifferentiated TYMS^+/+^- and TYMS^-/-^ -hiPSC aggregates subcutaneously in mice, without or with 100 mg/kg/day dTMP supplementation in the drinking water (Fig. 2.D). After 6 weeks, we retrieved the teratomas from the euthanized mice and analyzed their size, composition, and proliferation (Fig. 2.E-K).

*TYMS*^+/+^-hiPSC produced teratoma-like growth in all mice regardless of the thymidine supplementation, while *TYMS^-/-^*-hiPSC produced visible teratomas only under dTMP supplementation. Only one mouse presented a small *TYMS^-/-^*-derived teratoma without supplementation (Fig. 2.E). Histological analysis and immunocytochemistry showed that *TYMS*^+/+^-hiPSC generated dense teratomas with higher cellularity (Fig. 2.F-G). Contrarily, *TYMS*^-/*-*^-hiPSC overall produced low-density teratomas. As expected, immunohistochemical analysis of *TYMS*^-/-^ -hiPSC derived tumors resulted negative for TYMS, while *TYMS*^+/+^-hiPSCs present a rather homogenous expression of the protein (Fig. 2.H). Additionally, analogous to the *in vitro* experiment, we could identify regions expressing markers for the three germ layers (Fig. 2.I) in all analyzed tumors. Importantly, the absence of thymidine supplementation resulted in significantly less proliferative tumors with low cellularity, (Fig. 2.J-K), and composed mainly of primitive glial, epithelial, and mesenchymal tissue.

### Non-supplemented TYMS^-/-^ -hiPSC differentiate into functional beta cells *in vitro*

One promising target for cell replacement therapy is diabetes, in which stem cell derived insulin-producing beta cells act as the therapeutic source of insulin. We evaluated the safety, quality, and function of *TYMS*^-/-^ -hiPSC-derived pancreatic islets using a well-defined, multi-step protocol for beta cell differentiation (Fig. 3.A) (*3*, *4*). We differentiated *TYMS*^-/-^ -hiPSCs (HEL24.3) *in vitro*, supplemented with thymidine (40 µM) through the whole differentiation experiment. Meanwhile, as expected, non-supplemented *TYMS*^-/-^ cells did not survive thymidine withdrawal at early stages of differentiation, making further characterization of this condition impossible. Non-supplemented *TYMS*^+/+^-hiPSC were used as controls. We did not observe differences between *TYMS*^+/+^-hiPSC differentiated with or without dTMP supplementation, suggesting that the addition of dTMP at our working concentrations (up to 40 µM) does not affect *TYMS*^+/+^-cells (data not shown). Hereafter, we did not supplement *TYMS*^+/+^-hiPSC during the differentiation.

**Figure 3.**
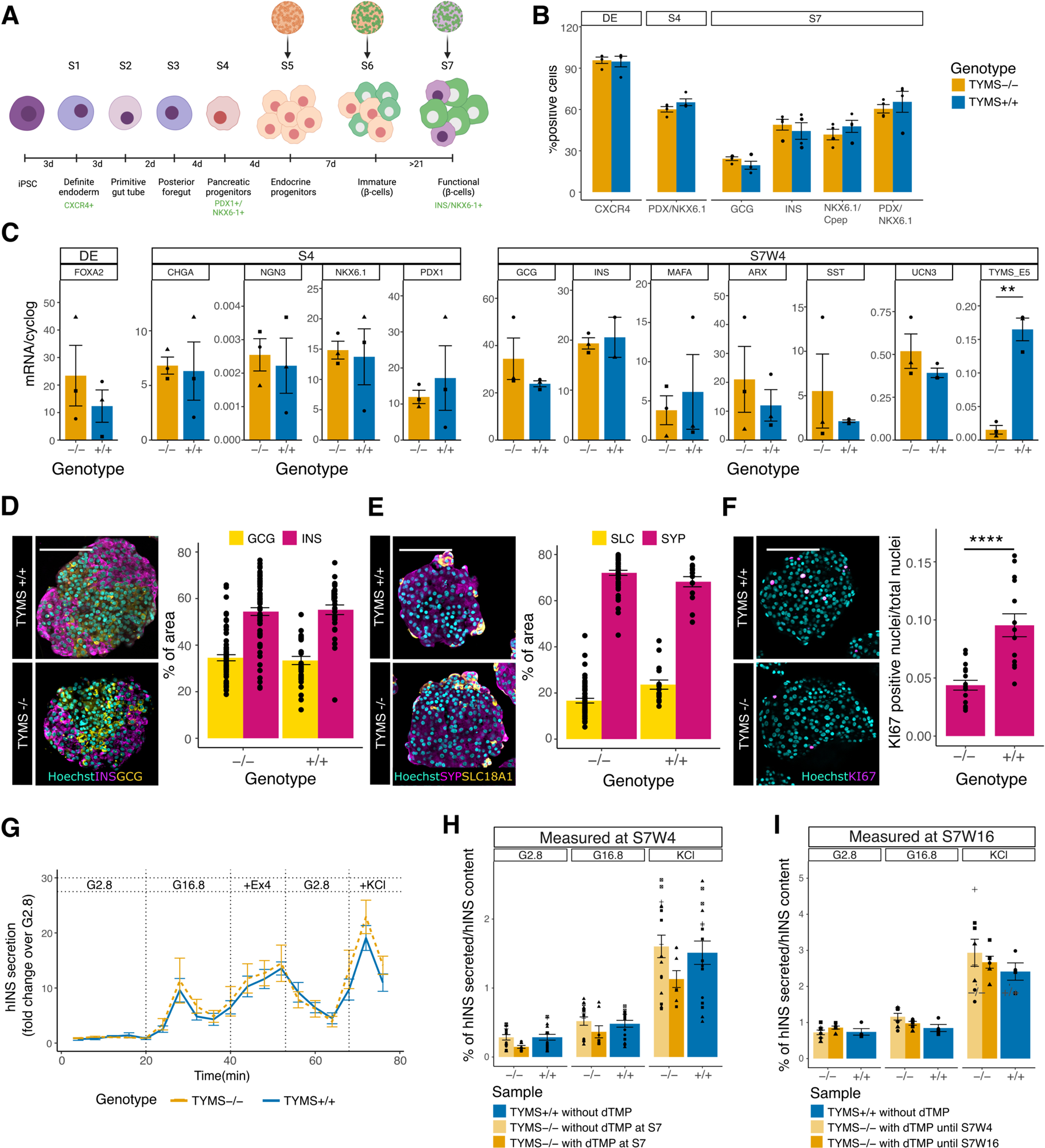
**A.** Graphical representation of in vitro beta cell differentiation from hiPSC. Most tests were carried out four weeks from stage 7 (S7W4), after four weeks from dTMP withdrawal. **B.** Flow cytometry analysis for markers for different stages of differentiation: CXCR4 for definite endoderm (DE) stage; NKX6.1 and PDX1 for stage 4 (S4); NKX6.1, C-peptide, PDX1, insulin (INS) and glucagon (GCG) for S7. **C.** mRNA expression analysis of maturation markers at different stages and of *TYMS* at S7 (primers target exon 5). Results are shown as ratio of marker mRNA over *GAPDH* mRNA. **D.** lmmunohistochemistry and quantification of INS (magenta) and GCG (yellow) in hiPSC-derive islets at S7W4. **E.** lmmunohistochemistry and quantification of synaptophysin (SYP, magenta) and SLC1SA1 (yellow) in hiPSC-derive islets at S7W4. **F.** lmmunohistochemistry of Kl67 (magenta) in hiPSC-derive islets at S7W4 and quantification of proliferation index, as number of Kl67 positive nuclei over the total nuclei. **G.** Dynamic insulin secretion in response to different stimuli at S7W4. **H-1.** In vitro insulin secretion in response to low (2.8) and high (16.8) glucose concentration and glucose plus KCI (2.8 KCI) at S7W4 (E) and S7W16 (F). Statistical significance based on Wilcoxon test; p>0.05 (ns, not shown), p < 0.05 (*), p < 0.01 (**), p < 0.0001 (****). Scalebar = 100 *µM*.

We evaluated the progression of the differentiation by flow cytometry, RT-qPCR, and immunocytochemical analysis of cell identity markers at several developmental stages. *TYMS*^+/+^- and supplemented *TYMS*^-/-^ -hiPS cells showed comparable *in vitro* expression levels of markers at the definitive endoderm (DE) stage (CXCR4), pancreatic endocrine progenitor stage (S4) (PDX1, NKX6.1) and endocrine maturation stage (S7) (insulin – INS, glucagon – GCG, NKX6.1, PDX1, C-peptide) by flow cytometry and immunocytochemistry (Fig. 3.B and Fig. S2.A-C). In transcriptomic analysis by RT-qPCR, we found that both the *TYMS*^-/-^ - and *TYMS*^+/+^-derived islets expressed similar levels of markers at stages DE and S4 stages, and markers of mature beta-cells at week 4 of stage 7 (S7W4) (Fig. 3.C). *TYMS*^-/-^ -derived islets presented lower *TYMS* mRNA levels that their *TYMS*^+/+^ counterparts. Hereafter, we withdrew dTMP supplementation from the islets cultured for *in vivo* implantation. By the last stage before implantation (S7W4), the *TYMS^-^*^/-^-hiPSC islets showed normal morphology and standard percentage of insulin, glucagon, synaptophysin (SYP) and SLC18A1 expressing cells (Fig. 3.D-E), while displaying a significantly decreased proliferative index by KI67 analysis (Fig. 3.F). Accordingly, *TYMS*^-/-^ and *TYMS*^+/+^ hiPSC-derived islets showed similar *in vitro* levels of glucose-, exendin 4- and KCl-induced functional insulin secretion at S7W4 (Fig. 4.G-E) and even after 3 months in culture (S7W16, Fig. 3.I). Thus, these results overall indicate that *TYMS*^-/-^ -hiPSC lines efficiently differentiate to functional SC-islets, comparable to their control counterparts, while displaying reduced proliferation upon dTMP withdrawal.

**Figure 4.**
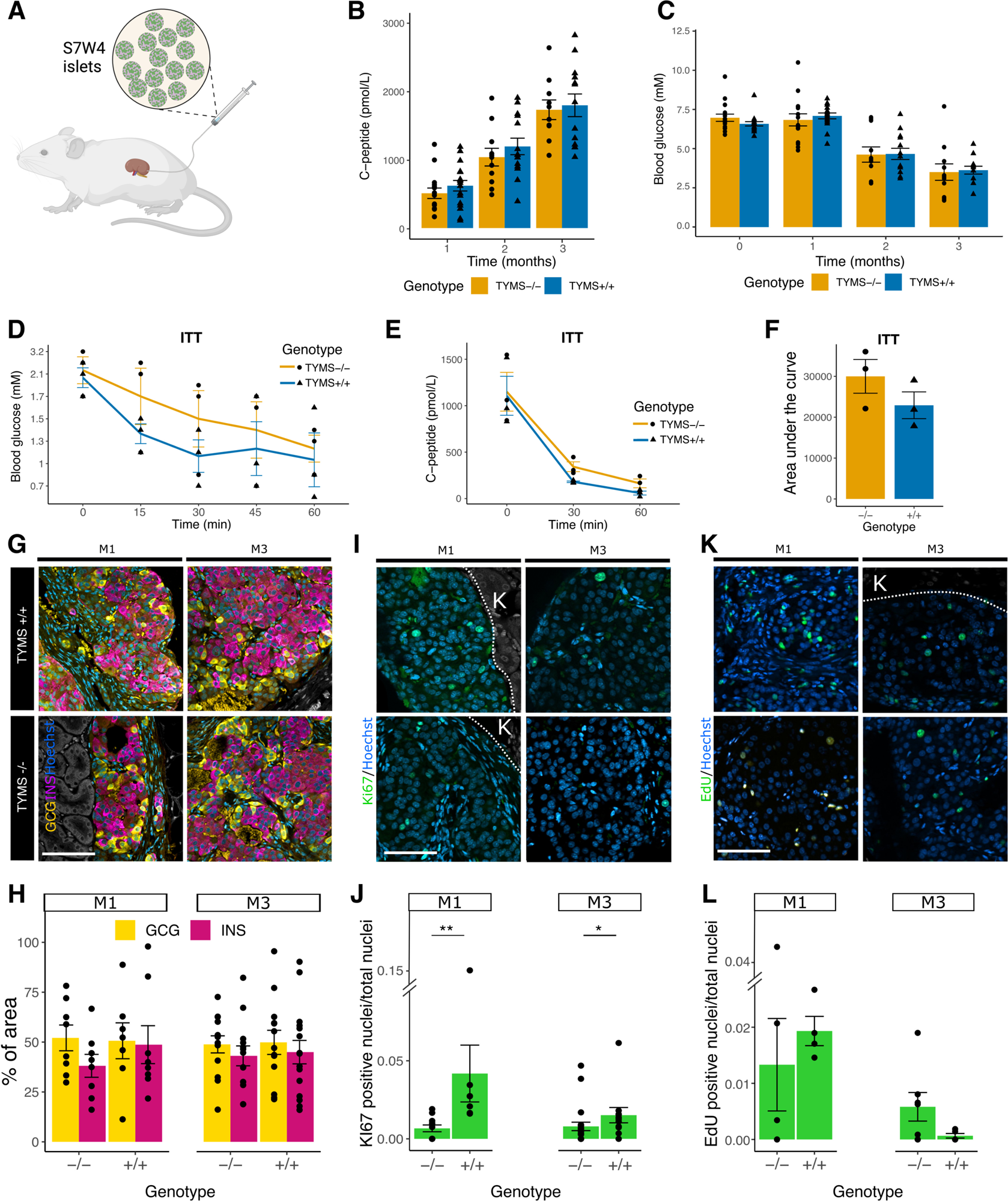
**A.** Diagram of S7W4 islet implantation set-up. **B-C.** Follow up of human C-peptide secretion (in serum), and blood glucose in mice after 1-, 2- and 3-months from implantation (month 0). **D.** Blood glucose of mice under ITT at every 15 min for 1 hour. **E.** Human C-peptide secretion (in serum) during ITT before and after 30 min and 1 hour from insulin administration. **F.** Quantification of area under the curves of human-C­ peptide secretion. **G.** lmmunohistochemistry of engrafted kidney 1- and 3-months post-implantation. INS (magenta), GCG (yellow). **H.** Quantification of INS and GCG area in the islets from G. I. lmmunohistochemistry against Kl67 in grafts after 1 and 3 months. **J.** Proliferation index, based on the quantification of Kl67 positive nuclei over the total number of nuclei from I. **K.** lmmunohistochemistry against EdU in grafts after 1 and 3 months. **L.** Quantification of EdU positive nuclei over the total number of nuclei from K. Statistical significance based on Wilcoxon test; p>0.05 (ns, not shown) p < 0.05 (*), p < 0.01 (**). K = kidney. Scalebar = 50 µM.

### *TYMS*^-/-^ -hiPSC derived beta cells without dTMP supplementation efficiently regulate blood glucose *in vivo*

We next evaluated the function of mature *TYMS*^-/-^ beta cells (S7) *in vivo* by engrafting them under the kidney capsule of immunosuppressed mice without dTMP supplementation (Fig. 4.A). We implanted mice with *TYMS*^+/+^- or *TYMS*^-/-^ -hiPSC -derived islets, and monitored their body weight, glucose levels and the grafts’ production of human C-peptide for three months.

Both *TYMS*^-/-^ and *TYMS*^+/+^ control islet produced comparable levels of human C-peptide and lowered the mouse blood glucose (Fig. 4.B-C). All mice maintained their normal body weights (Fig. S2.F) and did not show any other signs of health concerns.

Three months post-implantation, we performed an insulin tolerance test by treating the mice with insulin to test the ability of the grafts to shut down their *bona fide* insulin secretion in response to low glucose levels. Upon the decline of circulating glucose levels in response to the exogenously injected insulin (Fig. 4.D), our *TYMS*^-/-^ grafts stopped their insulin secretion at comparable rates as their *TYMS*^+/+^ counterparts, suggesting an appropriate regulation of this process (Fig. 4.E-F).

Immunohistochemistry on the implanted grafts showed comparable percentage of insulin and glucagon producing cells in *TYMS*^+/+^ and *TYMS*^-/-^ grafts at 1- and 3-months post-implantation (Fig. 4.G-H).

Additionally, to quantify the proliferation of the cells in the grafts *in vivo* through time (1 and 3 months), we ran immunohistochemical analysis against KI67 and we supplemented the drinking water of the mice with EdU for 5 days before euthanizing them. Interestingly, despite the significant reduction on active proliferation in *TYMS*^-/-^ cells (by KI67 expression analysis), the uptake of EdU did not differ significantly (Fig. 4.I-L and Fig. S3.D-E), possibly due to the uptake of EdU as a source of thymidine.

To summarize, we demonstrate here that *TYMS*^-/-^ -hiPSCs differentiate into pancreatic beta cells and function normally *in vitro* and *in vivo*, as seen by prolonged regulated human insulin secretion.

## Discussion

We have developed a method to generate controllable cells without inserting any external genetic elements. By disrupting the rate-limiting DNA-specific nucleotide synthesis reaction, we have obtained cells auxotrophic to thymidine. This novel safety mechanism allows *TYMS*^-/-^ -hiPSCs to expand as needed for undefined periods showing no signs of exhaustion, but only when externally supplemented with thymidine. In its absence, contrarily, the cells fail to proliferate. Moreover, this method does not affect function of the therapeutic differentiated cells *in vitro* or *in vivo*.

The *TYMS*^-/-^ -hiPSCs retain their pluripotency and differentiate successfully into different cell types. Non-supplemented *TYMS*^-/-^ -hiPSC generate small or no teratomas *in vivo* with a significantly lower proliferative index (KI67), pointing to the loss of proliferative capacity of these cells under physiological concentrations of thymidine. Previous reports by Diehl et al (*25*) suggest that thymidine withdrawal from *TYMS*^-/-^ hiPSCs leads to replicative stress and cell cycle arrest from nucleotide starvation. Supported by the results of our cell cycle analysis, we hypothesize that cells that have not exited the cell cycle die likely soon after this replicative stress is sensed, at the S-G2 checkpoint (*25*, *26*). However, we observed an extended survival time of non-supplemented *TYMS*^-/-^ cells when they form 3D aggregates both in *in vitro* and *in vivo* differentiation experiments (embryoid bodies and teratoma formation, respectively). As TYMS activity is required for the synthesis of DNA but not that of RNA, this may suggest the recycling of thymidine from apoptotic cells until cell cycle exit (*27*). Once they exit the cell cycle, the cells may survive as differentiated cells for extended periods. Accordingly, we see no functional alterations in the terminally differentiated cells without dTMP supplementation *in vitro* or *in vivo*. The hiPSC-derived pancreatic islets survived implantation and proved to successfully secrete human insulin and lower blood glucose up to the measured three months after implantation, without affecting the overall wellbeing of the mice.

The advances in basic science and technological developments simultaneously aid and pressure pre-clinical studies in their translation to clinical applications. On one hand, improved gene and cell manipulation protocols have increased the efficiency and widen the application of somatic-cell and gene therapies, tissue-engineered medicines, and CRISPR-based products. On the other hand, the safety concerns arising from altering DNA challenge further clinical applications. Hence, strategies to make cellular products safer become a need. Ideally, a safety mechanism for cell therapies selectively targets cells that retain or have acquired the ability for dangerous proliferation, without affecting the functional therapeutic cells. Although attempts to control the activity and proliferation of the therapeutic cells have proven relatively efficient, they present some inherent disadvantages. As most *safety switches* alter DNA by including exogenous genetic material, they often result complex, immunogenic or susceptible to mutations or silencing, ultimately affecting the partial/remissive function of the system (*20*).

In this regard, we believe our system brings added value to the efficiency and versatility of safety mechanisms. By a simple genetic manipulation of endogenous DNA, we generated cells auxotrophic towards thymidine. This approach not only evades the insertion of external genetic sequences, but additionally allows for control of cellular proliferation by addition of a simple, FDA-approved compound (thymidine). Thus, it facilitates the mass expansion and generation of therapeutic cells, that lose their proliferative capacity, but not their function, under *in vivo* concentrations of thymidine. Moreover, *in vivo* thymidine administration may control the function and efficiency of some therapeutic cells, such as immunotherapies, expanding the applications of this strategy.

The genetic disruption applied in this research brings about an additional advantage: the use of the *TYMS* locus as a genomic *safe harbor*. This opens the doors to future additions and combinatory developments with other selection and safety systems. We see this versatile feature as a solution to current limitations and an aid to expand the applications of *TYMS* disruption to other aims and other therapeutic cell types. We expect the replication and utilization of these results to contribute to the optimization of hiPSC-derived and other cellular products, and to the future development of safer efficient cell-based therapies.

## Aknowledgements

We thank MD Vaino Lithovius and MSc. Nidhi Madhusudan for assistance in image analysis, MSc. Sami Jalil, PhD. Diego Balboa and PhD. Juan Cruz Landoni for scientific discussion and support, MSc. Hazem Ibrahim and MSc. Eliisa Vahankangas for assistance with animal experiments. We thank the Flow cytometry Unit and the Animal facility at Biomedicum Helsinki for their possible contribution to this study.

We thank Helsinki University Hospital research funding, Foundation for Pediatric research, Paulo Foundation, Signe and Ane Gyllenberg Foundation, Magnus Ehrnrooth Foundation, K. Albin Johansson Foundation, Ida Montin Foundation, Finnish Red Cross Blood Service Research Fund, Diabetes Foundation for financial support.

## Author contributions

Conceptualization: RSM, TO, KW

Methodology: RSM, HM, JSV, KW

Investigation: RSM, HM, SE, IS, MB, JSV.

Visualization: RSM

Formal analysis: RSM

Funding acquisition: RSM, TO, KW

Project administration: RSM, KW

Supervision: TO, JSV, KW

Writing – original draft: RSM, KW

Writing – review & editing: RSM, HM, JSV, HP, TO, KW

## Materials and methods

### Culture of HEK293 FT, A431, A549, HSC3

We cultured HEK293, epidermoid carcinoma cell line A431, adenocarcinoma human alveolar basal epithelial cells A549, and human oral squamous carcinoma cell line HSC-3 cells in Dulbecco’s modified Eagle’s medium (DMEM) containing 10% fetal bovine serum (FBS), 100 µg/ml penicillin-streptomycin and 2 mM GlutaMAX (all ThermoFisher Scientific). After *TYMS* knock-out, cell media were additionally supplemented with 5-20µM dTMP (ThermoFisher Scientific). We kept the cell lines at 37 °C and 5% CO2, with media change every other day until splitting with TrypLE select (Gibco). All cell lines tested negative for mycoplasma.

### Induced Pluripotent Stem cells culture and characterization

Human induced pluripotent stem cell (hiPSC) line HEL24.3was cultured on Matrigel (Corning) coated plates in E8 medium (ThermoFisher Scientific) and split using 0.5 mM EDTA. We kept the cell lines at 37 °C and 5% CO2, with media change every other day. All cell lines tested negative for mycoplasma. After *TYMS* knock-out, cell media were additionally supplemented with 5-20 µM dTMP. The karyotyping was carried out by Ambar Anàlisis Mèdiques (Barcelona, Spain) by G-banding. We treated edited (passage 20 and 45) and non-edited (passage 20) hiPSC with colcemid for 4 h, 37°C, 5% CO2 and proceeded to prepare the cells as recommended by the service provider.

### Genome editing

To knock out exon 5, we designed guide RNA targeting intron 4 (I4) and intron 5 (I5) using online tools (https://benchling.com, CRISPOR (*28*), Table S1). We generated the sgRNA by incubating 5 min at 95 °C our gRNA (customized Alt-R CRISPR-Cas9 gRNA, iDT) with Alt-R® CRISPR-Cas9 tracrRNA, ATTO™ 550 (iDT).

For each stem cell electroporation experiment, we dissociated 2×10^6 cells into single-cells with StemPro Accutase (ThermoFisher Scientific). We complexed the sgAlt-R® S.p. HiFi Cas9 Nuclease V3 and and both sgRNA to form the functional ribonucleoprotein (RNP) and delivered this RNP into the cells, along the Alt-R® Cas9 electroporation enhancer (all from integrated DNA Technologies – iDT), by electroporation with Neon transfection systems (1100 V, 20 ms, 2 pulses). Cells were plated onto Matrigel-coated plates containing E8 with 5 µM ROCK inhibitor (Y-27632, Selleckchem) and 20-40 µM dTMP, and incubated at 37 °C, 5% CO2.

For electroporation experiment with cell lines HSC-3 and A549, A431, cells were dissociated into single cells with TrypLE. SgRNAs were prepared as previously described and delivered to the cells using Neon transfection systems. For HSC3 and A549, the electroporation settings were 1250 V, 10 ms, 1 pulse, while for A431 we used 1450 V, 20 ms, 2 pulses. After tranfection, the cells were plated onto plates containing culture media (10% FBS in DMEM) supplemented with 20-40 µM dTMP, and incubated at 37 °C, 5% CO2.

24-48 h after electroporation, we single cell sorted ATTO550 positive cells for monoclonal expansion in 96-well plates containing their corresponding dTMP-supplemented culture media (hiPSC additionally supplemented with 10% Clone R (StemCell Technologies)). The media was refreshed every 72 h until splitting. We individually screened monoclonal colonies by PCR. All PCR products of the selected edited clones were validated by Sanger sequencing, along with the integrity of the top-7 off target sequences (Table S2).

### Embryoid body differentiation

To test the pluripotency for an embryonal lineage differentiation, we grew the cells until 90-100% confluency and performed embryoid body (EB) differentiation assay as previously described (*2*). Final plated EBs were fixed with 4%PFA-PBS and analyzed by immunocytochemistry for characteristic markers of the three germ-layers (see list below).

### Teratoma assay

Knock-out and wild-type HEL24.3 cells were cultured in ultra-low attachment plates with E8 supplemented with 5 µM ROCK inhibitor for 24-48 h to form aggregates. The medium for knock-out cells was additionally supplemented with 40 µM dTMP until 2 h before implantation. Groups of approximately 500 spheres were collected in syringe cannulas for subcutaneous implantation in the legs of the mice. For half of the mice, the drinking water was supplemented with 100 mg/kg/day dTMP. 6-8 weeks post implantation the teratomas were extracted and fixed for immunocytochemistry and hematoxilin-eosin stainings as described below.

### dTMP auxotrophy

Wild-type and knock-out clones from each cell line were split onto Matrigel coated 6-well plates at a reason of a 25000 cells per well. Each well contained the corresponding medium for the cell line (see *Cell culture*) supplemented with 0 to 5 mM dTMP. Cells were washed and media were refreshed daily to clear away dead cells debris and keep the thymidine concentration as constant as possible.

### Beta-cell differentiation

The *in vitro* beta cell differentiation was carried out as previously described (*3*, *4*). Briefly, *TYMS* knock-out and control HEL24.3 hiPS cells were seeded onto Matrigel coated 10 cm plates as previously stated. Differentiation protocol was started 24 h post-seeding, when the medium was changed to D0 medium. Both cell lines were supplemented with 0 or 40 µM dTMP until maturation. At stage 7, dTMP was withdrawn or kept until analysis (weeks 4 to 16 from the first day of S7). The media was refreshed every 2-3 days until analysis.

### Flow cytometry

Stage 4 cells and stage 7 stem cell derived islets (SC-islets) were dissociated with TrypLE for 10 min at 37 °C and resuspended in 5% FBS-containing PBS. Fixation and permeabilization were done using Cytofix/Cytoperm (BD Biosciences, catalog no. 554714) for 20 min at RT. Then, samples were incubated overnight with primary antibodies at 4 °C, followed by secondary antibodies for 30 min in RT in Perm/Wash buffer (BD Biosciences, catalog no. 554714) supplemented with 5% FBS. The cells were run on FACSCalibur cytometer (BD Biosciences); data were collected with CellQuest Pro v.4.0.2 (BD Biosciences) and analyzed with FlowJo™ v10.8 Software (BD Life Sciences). Antibodies are listed in Table S3.

For cell cycle analysis, wild type and knock out hiPSC were plated overnight on 6 well plates containing E8 supplemented with ROCK inhibitor and 20 µM dTMP. Then, the media were refreshed and for some plates the dTMP supplementation was stopped. Cells from individual wells were collected at 0, 2, 4, 10, 16 and 24 h after dTMP withdrawal and fixed using 4% PFA in in PBS. The same timepoint were collected for the supplemented plates. After fixation, samples were washed and incubated 30 min with 10 μg/ml DAPI diluted in 1% Triton-X (Sigma-Aldrich) in PBS. Then, they were immediately analyzed using the NovoCyte Quanteon 4025.

### Insulin secretion analysis *in vitro*

For static analysis of insulin secretion, a total of 30 – 50 SC-islets were picked and incubated for 90 min in a 12 well plate containing 2.8 mM glucose (G2.8) in Krebs-Ringer buffer (KRB) for equilibration. This was followed by sequential 30 min incubations of G2.8, 16.8 mM glucose (G16.8) and G2.8+30 mM KCl in KRB. After each incubation, 200 uL samples were taken for further insulin secretion analysis. Once all samples were taken, the islets were collected for DNA quantification. Dynamic insulin secretion test was carried out using a perifusion apparatus (Brandel Suprafusion SF-06) at a 0.25 mlmin–1 flow rate, sampling every 4 min. 50 SC-islets were handpicked and perfused with KRB; sample collection started after 90 min of equilibration in G2.8. The insulin content of secretion fractions and SC-islet lysates was analyzed with enzymelinked immunosorbent assay (ELISA) (Mercodia).

### Islet *in vivo* characterization

Animal care and experiments were approved by the National Animal Experiment Board in Finland (ESAVI/9734/2021). NOD-SCID-Gamma (NSG, Jackson Laboratories, catalog no. 0055577) mice were housed in the Biomedicum Helsinki conventional facility in 12 h light/dark cycle and fed standard chow. SC-islet implantations were done following a previously stablished described protocol (*4*). Non-fasted blood samples were collected monthly from the saphenous vein for glucose measurement and C-peptide secretion analysis. For further characterization of the SC-islet graft, the engrafted kidney was removed after 1 or 3 months and processed as described below.

For the insulin tolerance test at 3-months post-implantation, the mice were weighted and injected with insulin (0.75 IU/kg). Blood glucose was measured every 15 min for 1 hour and serum samples for human C-peptide secretion analysis were at 30 and 60 min.

### Hematoxylin-eosin staining

Teratoma-like growths were fixed using 4% PFA-PBS overnight and then embedded in paraffin and sectioned.

Dried sections were deparaffinized and stained using a standard protocol. Briefly, they were deparaffinized using xylene (3 x 10 min) and gradually hydrated using decreasing concentrations of ethanol (99% EtOH 3 x 4 min, 96% EtOH 1 x 4 min, 70% EtOH 1 x 2 min). Then incubated in hematoxylin for 2.5 – 3 min, washed in indirect flowing tap water for 5 - 10 min followed by a 1 min incubation in MQ water. The sample were incubated in eosin for 2 min and dehydrated using increasing concentrations of ethanol (96% EtOH 3 x 10 sec, 99% EtOH 2 x 2 min) and xylene (2 x 4min). Slides were mounted with coverslips using mounting media and image the next day.

### Immunocytochemistry, immunohistochemistry, and image analysis

Samples from hiPSC were fixed using 4%PFA-PBS for 20 min and permeabilized using 1% Triton-X in PBS. Samples from S4 and S7 SC-islets were fixed with 4% PFA in PBS for 2 h, and explanted SC-islet grafts and teratoma-like growths were fixed overnight. After fixation, these samples were embedded in paraffin and sectioned.

For immunohistochemistry and immunocytochemistry, 5 μm sections were deparaffinized and subjected to HIER in 0.1 mmol l–1 citrate buffer. The cells and tissue slides were blocked with UV-block (Thermo Scientific, catalog no. TA-125-PBQ), and incubated with primary antibodies in 0.1% Tween-20 overnight at 4 °C with the given dilutions (Table S3). After washing, secondary antibodies diluted in a similar manner were incubated at RT for 1 h. For immunocytochemistry, secondary antibody incubation was done in the presence of Hoechst33342 for nuclear staining. For immunohistochemistry, after the secondary antibody the slides were incubated with a DAB solution for 2-10 min until a brown color developed. Then, they were counterstained with hematoxylin for 2.5-3 min and dehydrated following a standard protocol (see hematoxylin-eosin staining).

The cells and immunocytochemistry slides were imaged using Apotome II from a Zeiss AxioImager with the same exposure and export setting used on all slides of each targeted marker. Immunohistochemisty slides were imaged using an Olympus BX51 Microscope.

Images were processed in Zen2 Blue Edition v.2 (Zeiss) and analyzed using FiJi (*29*) or CellProfiler v.4.0 (*30*) with pipelines adapted from a previous report (*31*). The same pipeline settings were used on all images of each targeted marker.

### Protein expression analysis

mRNA expression levels were analyzed by qPCR (Table S4-6). TYMS protein expression was assessed by standard Western blot. Briefly, protein was extracted by using RIPA lysis buffer (20-188, Millipore) supplemented with cOmplete Mini protease and phosphatase inhibitors tablets (Roche). Samples were sonicated (30 sec, %50 duty cycle pulse) and centrifuged for soluble protein extraction. Protein quantification was done using a Pierce BCA protein assay kit (ThermoFisher) and 20-30 μg total protein were run in Mini-PROTEAN TGX Precast gels (#4561033, Bio-Rad). Transfer to a nitrocellulose membrane was done using 25 V for 6 min in the iBlot2 Gel Transfer device (IB21001, ThermoFisher). Then, the membrane was probed with a rabbit polyclonal primary antibody against TYMS (1:2500, Proteintech; Table S3). For detection, we used an anti-rabbit HRP-coupled secondary antibody (1:5000, Cell Signalling) and Western ECL substrate (#1705061, Bio-Rad. As a loading control, we probed the membrane with a b-actin-HRP coupled antibody (sc-47778, Santa Cruz).

---

**Supplementary figure 1.**
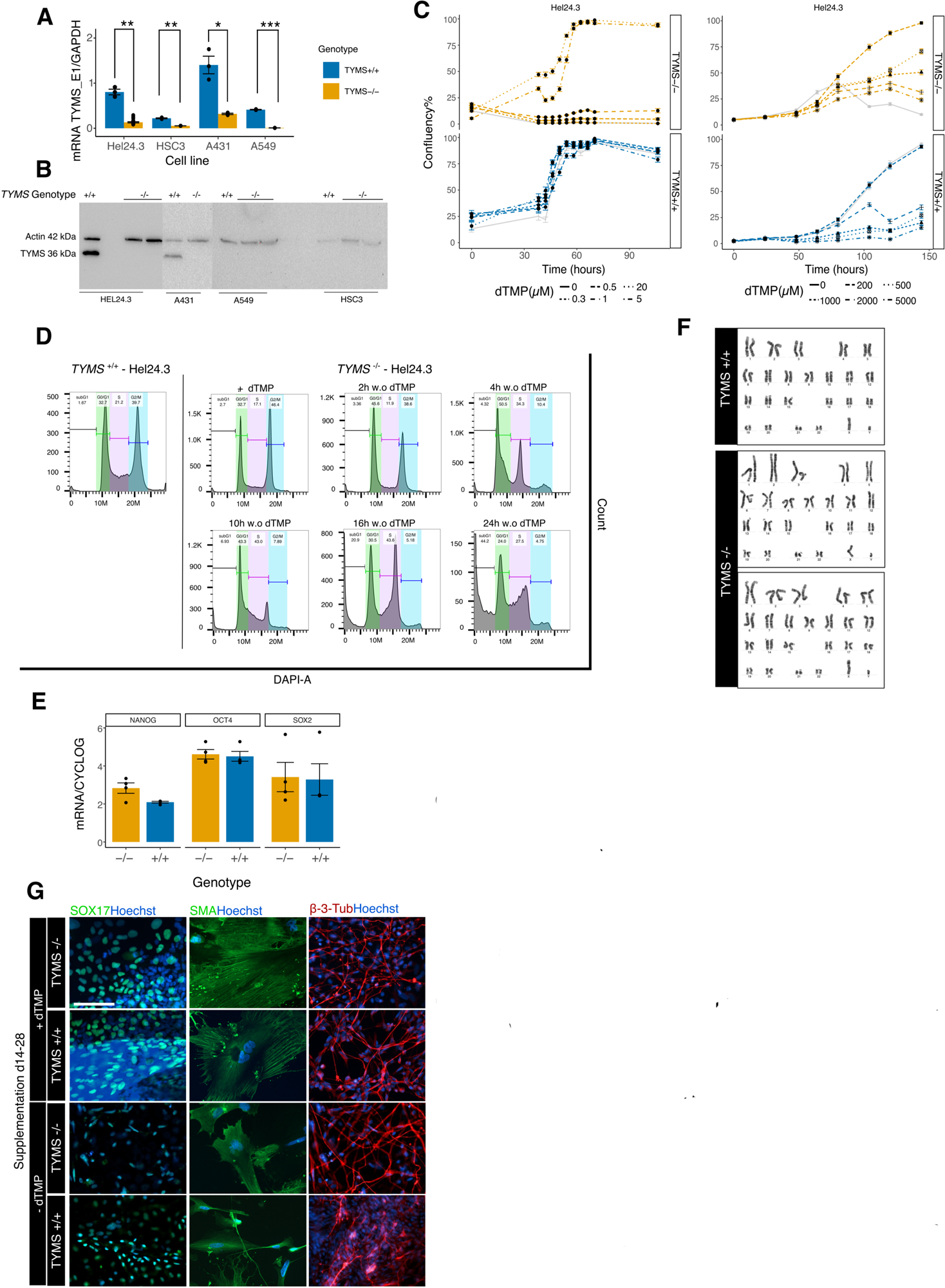
**A.** *TYMS* mRNA expression in wild-type versus knock-out cell lines (HEL24.3, A431, A549 and HSC3) using primers targeting exon 1 (n=3/cell line). Results shown as ratio of *TYMS* mRNA over *GAPDH* mRNA. **B.** Full western blot membranes of TYMS in wild-type and edited cell lines. **C.** Full growth curves of wild-type and knock-out hiPSC under different concentrations of dTMP (0 to 5000 µM). Results shown as average confluency per image ± SD. **D.** Cell cycle analysis of knock-out hiPSC during the first 24h after dTMP withdrawal. **E.** Karyotype of wild-type hiPSC, passage 20. Karyotype of TYMS-knock-out hiPSC, passage 20 (up) and passage 45 (down). **F.** hiPSC RT­ qPCR for expression analysis of pluripotency markers *NANOG, OCT4* and *SOX2.* Results shown as ratio of mRNA over *CYCLOG* mRNA. **G.** lmmunocytochemistry against markers for endoderm (SOX17), mesoderm (SMA), and ectoderm (beta-3-tubulin) in cells derived from wild-type and knock-out hiPSC without dTMP supplementation during the first stage of differentiation. Statistical significance in panel A and F based on Wilcoxon test; p < 0.05 (*), p < 0.01 (**), p < 0.001. Scalebar = 100 µM.

**Supplementary figure 2.**
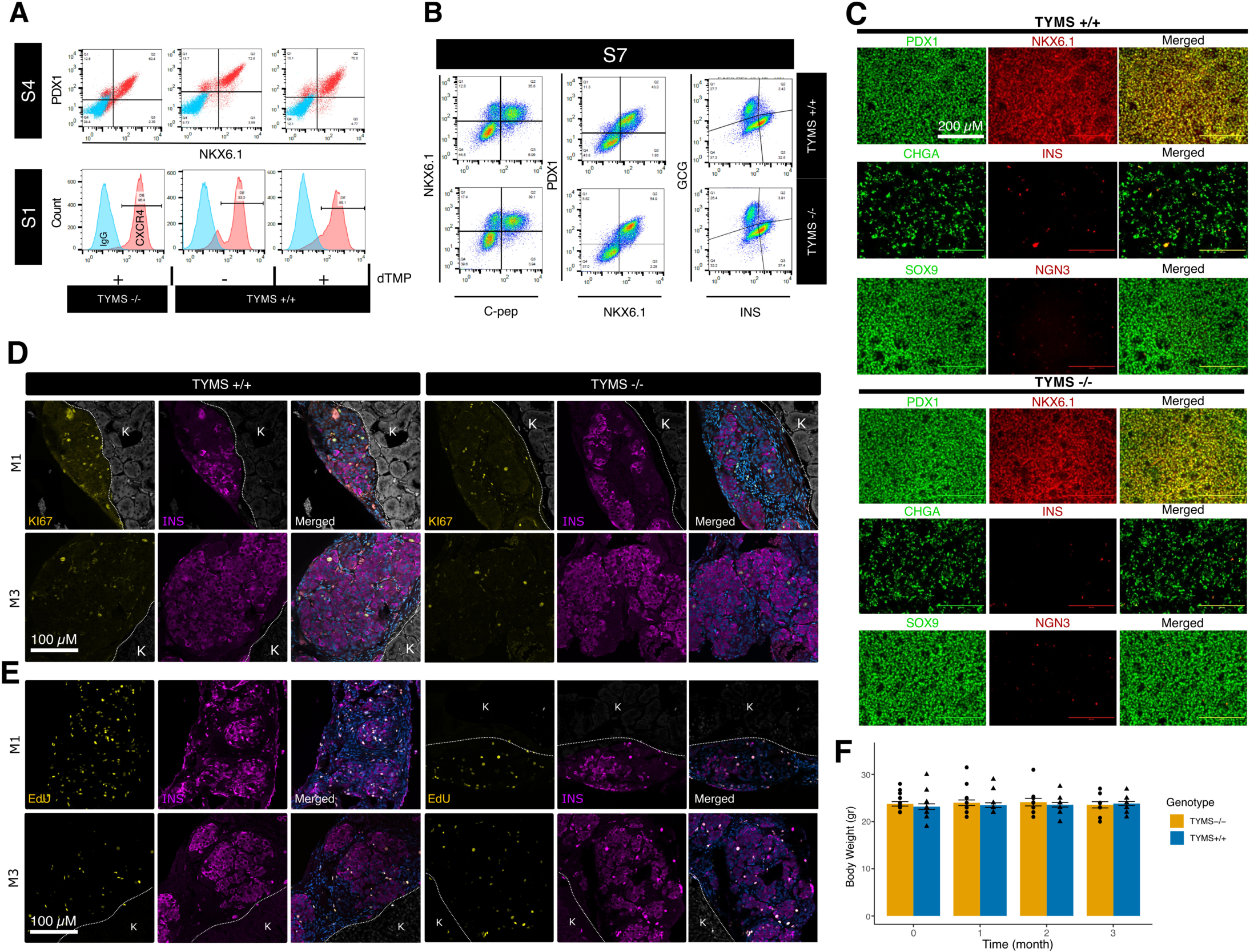
**A.** Flow cytometry analysis of characteristic markers for definite endoderm (CXCR4) and stage 4 (NKX6.1 and PDX1) of beta-cell differentiation protocol. **B.** Flow cytometry analysis of markers at stage 7 of beta-cell differentiation: NKX6.1, C-peptide, PDX1, insulin (INS) and glucagon (GCG). **C.** lmmunocytochemistry analysis of maturation markers at S4 of beta-cell differentiation protocol. First row: PDX1 and NKX6.1. Second row: CHGA and INS. Third row: SOX9 and NGN3. **D-E.** lmmunocytochemistry against Kl67 (yellow, D) and EdU (yellow, E) on insulin positive cells (magenta) from beta-cell grafts at 1 or 3 months after implantation under the kidney capsule (M1 and M3, respectively). **F.** Body weight of mice after 1-, 2- and 3-months from implantation (month 0). K = kidney

