## Supplementary Tables for "Thymidylate synthase disruption to limit cell proliferation in cell therapies"

Table 1 – Summary table of selected “safety” systems, and potential advantages and disadvantages.

| **Type** | **Function** | **Examples** | **Advantages** | **Disadvantages** |
| --- | --- | --- | --- | --- |
| Enzyme/Prodrug  (*20*, *32*–*36*) | Suicide transgenes into cells that transform prodrug into toxic metabolite | HSV-TK + Ganciclovir (*37*, *38*) | Some have the ability to eliminate the whole graft  Could be combined with other strategies  Safe  Generally efficient | Transgenes derive from viruses, bacteria or yeast  Immuno-rejection of the therapeutic cells  Toxicity of the prodrug  Bystander effect  Slow activation time  Silencing of transgene |
|  |  | CD+5-FC (*39*) |  |  |
|  |  | NTR+CB1954 (*40*) |  |  |
|  |  | PNP+MEP (*41*) |  |  |
|  |  | mTMPK+AZT (*42*) |  |  |
| MAb mediated (*35*, *36*) | Engineer a gene targeted by an Ab into the transplanted cells | CD20t+Rituximab | Avoid GVHD (*43*)  Efficient  Different payloads for ADC | CD20 present in endogenous B cells |
|  |  | hEGFRt+Cetuximab (*44*) |  |  |
|  |  | c-myc tag + Anti-c-myc tag (*45*) |  |  |
|  |  | X+ADC anti-X (*46*) |  |  |
| Inducible Dimerization (*35*) | Insertion of a modified Caspase-9. The recruitment sequence is replaced by a dimerization domain of FK506 binding protein | iCas9 (*47*)+CID | Low immunogenicity  Inert small molecule  Fast action  Tested to eliminate iPSC (*48*) | Silencing of transgene  CID resistance (*49*) |
| Metabolic | Endogenous disruption of pyrimidine de novo synthesis | UMPS KO (*15*) | Transgene-free | May affect RNA synthesis |
| CID: Chemical inducer of dimerization; ADC: antibody–drug conjugate; GVDH: Graft-versus-host disease; CD: cytosine deaminase; NTR: Nitroreductase; HSV-TK: herpes simplex virus thymidine kinase; PNP: purine nucleoside phosphorylase; mTMPK: mutated thymidylate monophosphate kinase; OxPD: Oxazaphosphorine drugs; AZT: azidothymidine | | | | |

Supplementary table 1 – gRNAs used for CRISPR-Cas9 *TYMS* knock-out

| **Gene** | **Target** | **gRNA sequence** | **PAM** | **On target** | **Off target** |
| --- | --- | --- | --- | --- | --- |
| TYMS | Intron 4 | CAACTCATATGGTGGAGACC | AGG | 94 | 54 |
|  | Intron 5 | TCTGTTAGTGCGTATACCAC | AGG | 77 | 88 |

Supplementary table 2 – PCR and sequencing primer sequences for CRISPR-Cas9 off-targets

| **gRNA** | **Location** | **MM** | **Primer Sequence (5’-3’)** | **Product length** | **Algorithm** | **Assay** |
| --- | --- | --- | --- | --- | --- | --- |
| 1 | chr5_54455571 | 4 | AAAGTGCACTGCTGACTGGG | 586 | CRISPOR | PCR/S |
|  |  |  | AGACCACTTCTCAGAGGGGA |  |  |  |
|  | chr4_24900433 | 4 | CACCCACCATTCTGAGGACC | 528 |  | PCR/S |
|  |  |  | ATCCGTGTCACCATTCCCAC |  |  |  |
|  | chr8:-11058099 | 2 | TCCTGACTCCTTCAGTGGGG | 404 | Benchling | PCR/S |
|  |  |  | AAAGATACCACCGCCTCCCA |  |  |  |
|  | chr17:-43138461 | 4 | GTCGGTCCCAGGTGTTTCTC | 587 |  | PCR/S |
|  |  |  | ACCACTGGCTTTCAGGCTAC |  |  |  |
|  | chr19:+7152901 | 2 | CATTCAGACTCCACCCACCC | 373 |  | PCR/S |
|  |  |  | TCAGCCGCAGAGACTTGAG |  |  |  |
|  | chr14:+44505493 | 4 | GAGGCTGAAGCTCAAGAGGG | 514 | iDT | PCR/S |
|  |  |  | GCCCCAGCTAAAGTAGAGCC |  |  |  |
| 2 | chr1_233467802 | 4 | AGGCCCTGTAACTCCCTTCT | 600 | CRISPOR | PCR/S |
|  |  |  | CCAGCACCATCACTCCAAGT |  |  |  |
|  | chr13_26897943 | 3 | TGCACGTTCAGCTTGTGACT | 507 |  | PCR/S |
|  |  |  | ACAACTCTGCCTCACATGGAG |  |  |  |
|  | chr1_38172195 | 4 | CCATCCGATTGTAGTAGGCCC | 596 |  | PCR/S |
|  |  |  | TCCAGCTGGGCAATACTGTG |  |  |  |
|  | chr2:+13066090 | 3 | TCCCCACCCTATCTACTACCTC | 428 | Benchling/iDT | PCR/S |
|  |  |  | CAAATTATCCTGGGAATAAATGCAC |  |  |  |
|  | chr18:-31163947 | 3 | AACACCCATGCTGCTGAGAA | 639 | Benchling | PCR/S |
|  |  |  | TGAGCAGTGCCTGGAATCTC |  |  |  |
|  | chr9:+86563560 | 4 | CATGAGGTGGCTCAGTGGAG | 509 | Benchling/iDT | PCR/S |
|  |  |  | CCATGGCTCCCAATGCAGTA |  |  |  |

Supplementary table 3 – Antibodies used for immunocytochemistry, immunohistochemistry western blot, and flow cytometry

| **Epitope** | **Origin animal** | **Conjugate** | **Dilution** | **Supplier** | **Assay** |
| --- | --- | --- | --- | --- | --- |
| Nanog | Rabbit | N/A | 1: 500 | Cell Signaling;  #D73G4 | ICC |
| OCT4 | Mouse | N/A | 1:500 | Santa Cruz;  #sc-8628 | ICC |
| Sox2 | Rabbit | N/A | 1:500 | Cell signalling;  #D6D9 | ICC |
| SSEA-4 | Mouse | N/A | 1:500 | ThermoFisher;  #MA1-023 | ICC |
| Beta-3-  tubulin | Rabbit | N/A | 1:500 | R&D Systems;  #MAB1195 | ICC |
| SMA | Mouse | N/A | 1:500 | Sigma-aldrich;  #A2547 | ICC |
| Sox17 | Goat | N/A | 1:500 | R&D Systems;  #AF1924 | ICC |
| Ki-67 | Rabbit | N/A | 1:500 | Leica Microsystems  #NCL-Ki67p | ICC/IHC |
| Insulin | Guinea pig | N/A | 1:500 | Dako  #A0564; | IHC/FC |
| Glucagon | Mouse | N/A | 1:500 | Sigma-Aldrich  #G2654 | IHC/FC |
| Syn | Rabbit | N/A | 1;250 | Novus Biologicals;  #NB120-16659 | ICC/IHC |
| NGN3 | Sheep | N/A | 1:500 | R&D Systems  #AF3444 | ICC/IHC/FC |
| PDX1 | Goat | N/A | 1:250 | R&D Systems  #AF2419 | ICC/IHC/FC |
| NKX6.1 | Mouse | N/A | 1:250 | DSHB  #F55A10 | ICC/IHC/FC |
| CHGA | Rabbit | N/A | 1:500 | Dako  #A0564 | ICC/IHC/FC |
| SOX9 | Rabbit | N/A | 1:500 | Millipore  #AB5535 | ICC/IHC/FC |
| TYMS | Rabbit | N/A | 1:2500 | Proteintech;  #15047-1-AP | WB |
| Guinea pig | Goat | Red 594 | 1;500 |  | ICC/IHC/FC |
| Rabbit | Donkey | Red 594 | 1:500 | Thermofisher:  #A21207 | ICC/IHC/FC |
| Rabbit | Donkey | Green 488 | 1:500 | Thermofisher:  #A21206 | ICC/IHC/FC |
| Mouse | Donkey | Green 488  Red 594 | 1:500 | Thermofisher:  #A21202  #A21203 | ICC/IHC/FC |
| Goat | Donkey | Green 488 | 1:500 | Thermofisher;  #A11055 | ICC |
| Rabbit | Goat | HRP | 1;5000 | Cell Signalling;  #7074S | WB |
| B-actin | Mouse | HRP | 1;5000 | Santa Cruz;  #sc-47778 | WB |
| FC = Flow cytometry ICC = Immunocytochemistry  IHC = Immunohistochemistry WB = Western blot | | | | | |

Supplementary table 4 – PCR, qPCR and Sanger sequencing primer sequences for TYMS

| **Gene** | **Target** | **Sequence (5’-3’)** | **Product length** | **Assay** |
| --- | --- | --- | --- | --- |
| TYMS | Intron 4 | TCAACTCTACCAGGGTGTAG | 1336(WT)  871(KO) | PCR/SS |
|  | Intron 5 | CCAACCTCAGCATAGCTTTTG |  |  |
|  | Exon 2 | CCTCTGCTGACAACCAAACG | 95 | qPCR |
|  | Exon 3 | GAAGACAGCTCTTTAGCATTTG |  | qPCR |
|  | Exon 4 | TCAGGACAGGGAGTTGACCA | 117 | qPCR |
|  | Exon 5 | CAGCGCCATCAGAGGAAGAT |  | qPCR |
| SS: Sanger Sequencing | | | | |

Supplementary table 5 – qPCR primer sequences for pancreatic differentiation markers

| **Process** | **Gene** | **Differentiation stage** | **Sequence (5’-3’)** | **Product length** |
| --- | --- | --- | --- | --- |
|  | GAPDH | Housekeeping | GGTCATCCATGACAACTTTGG | 84 |
|  |  |  | CCATCCACAGTCTTCTGGGT |  |
| Pancreatic differentiation | FOXA2 | Definite Endoderm (DE) | AAGACCTACAGGCGCAGCT | 93 |
|  |  |  | CATCTTGTTGGGGCTCTGC |  |
|  | **CHGA** | **4** | AACCGCAGAC CAGAGGACCA | 102 |
|  |  |  | GTCTCAGCCC CGCCGTAGT |  |
|  | NGN3 | 4 | GACGACGCGAAGCTCACCAA | 98 |
|  |  |  | TACAAGCTGTGGTCCGCTAT |  |
|  | NKX6.1 | 4 | TATTCGTTGGGGATGACAGAG | 91 |
|  |  |  | TGGCCATCTCGGCAGCGTG |  |
|  | PDX1 | 4 | AAGTCTACCAAAGCTCACGCG | 52 |
|  |  |  | CGTAGGCGCCGCCTGC |  |
|  | GCG | 7 | GAAGGCGAGATTTCCCAGAAG | 113 |
|  |  |  | CCTGGCGGCAAGATTATCAAG |  |
|  | INS | 7 | CAGAAGCGTGGCATTGTGGA | 82 |
|  |  |  | GCTGCGTCTAGTTGCAGTAG |  |
|  | MAFA | 7 | GCCAGGTGGAGCAGCTGAA | 77 |
|  |  |  | CTTCTCGTATTTCTCCTTGTAC |  |
|  | ARX | 7 | ACAGACGCGCCTCTAGCATA | 81 |
|  |  |  | GCAGGATGTTGAGCTGCGTG |  |
|  | SST | 7 | CCCAGACTCCGTCAGTTTCT | 88 |
|  |  |  | ACAGCAGCTCTGCCAAGAAG |  |
|  | UCN3 | 7 | GCCACAAGTTCATGGGGACGTG | 120 |
|  |  |  | GACCGGCATCAGCATCTCTCC |  |

Supplementary table 6 - qPCR primer sequences for pluripotency markers

|  | **Gene** | **Sequence (5’-3’)** | **Product length** |
| --- | --- | --- | --- |
| Pluripotency | OCT4 | TTGGGCTCGAGAAGGATGTG | 91 |
|  |  | TCCTCTCGTTGTGCATAGTCG |  |
|  | SOX2 | GCCCTGCAGTACAACTCCAT | 85 |
|  |  | TGCCCTGCTGCGAGTAGGA |  |
|  | NANOG | CTCAGCCTCCAGCAGATGC | 94 |
|  |  | TAGATTTCATTCTCTGGTTCTGG |  |
|  | Cyclog | TCTTGTCAATGGCCAACAGAG | 84 |
|  |  | GCCCATCTAAATGAGGAGTTG |  |
